# Backbone Thioamide Substitution Enhances the Activity of Short Peptides in Modulating the Aggregation of α-Synuclein

**DOI:** 10.64898/2026.08.25.746879

**Authors:** Haoliang Zheng, Kyren Miller, Magdalena I. Ivanova, Robert W. Newberry

**Affiliations:** Department of Chemistry, The University of Texas at Austin, Austin, TX, USA; Biophysics Program, University of Michigan, Ann Arbor, MI, USA; Michigan Neuroscience Institute, University of Michigan, Ann Arbor, MI, USA

## Abstract

The non-amyloid-β component (NAC) region of the Parkinson’s-associated protein α-synuclein plays a key role in its pathogenic aggregation, motivating the development of molecules that target this critical region. Here, we show that a minimal NAC-derived motif, ^66^VGGAVVT^72^, can be reprogrammed through backbone engineering to modulate α-synuclein aggregation. Backbone thioamide substitution of this peptide enhances its interactions with α-synuclein fibrils and accelerates aggregation, whereas *N*-methylation disrupts β-sheet hydrogen bonding and inhibits fibrillization. Strikingly, combining these modifications yields hybrid peptides that inhibit the fibrillization of full-length α-synuclein at sub-stoichiometric concentrations. Consistent with in vitro results, these backbone-modified peptides can also reduce seeded α-synuclein aggregation in cells. These results establish that minimal amyloidogenic sequences can be systematically tuned from aggregation promoters to inhibitors through backbone-level perturbations, particularly thioamide incorporation.

## Introduction

Misfolding and aggregation of α-synuclein into β-sheet-rich assemblies are central pathological features of Parkinson’s disease and related synucleinopathies (Polymeropoulos et al., 1997; Spillantini et al., 1997). Accumulation of α-synuclein aggregates in Lewy bodies is strongly associated with neuronal dysfunction and degeneration (Goedert et al., 2013; Spillantini et al., 1997; Volpicelli-Daley et al., 2011). Fibrillar α-synuclein assemblies in particular are capable of templating the aggregation of endogenous α-synuclein and propagating inclusion pathology and neuronal loss throughout connected brain regions (Luk et al., 2012; Volpicelli-Daley et al., 2011). The aggregation process is driven by sequence-specific intermolecular interactions within the non-amyloid-β component (NAC) region (Jakes et al., 1994; Uéda et al., 1993), a hydrophobic segment that forms the core of the amyloid structure. Structural and biochemical studies have identified multiple aggregation-prone motifs within this region (Giasson et al., 2001; Goldschmidt et al., 2010; Murray et al., 2003; Murray et al., 2022a), highlighting the NAC domain as a critical determinant of α-synuclein misfolding (El-Agnaf et al., 1998; Han et al., 1995) and a key target for therapeutic intervention (Buell et al., 2014; Rodriguez et al., 2015).

As part of ongoing efforts to combat protein misfolding (Chiti and Dobson, 2017; Murray et al., 2023), we sought to identify critical motifs within misfolded α-synuclein that could be targeted by inhibitor candidates. α-Synuclein forms structurally different fibrils in which the subunits adopt distinct conformations, known as polymorphs or strains, which can be associated with distinct synucleinopathies (Schweighauser et al., 2020; Yang et al., 2022). Upon examining of α-synuclein structures from fibrils derived ex-vivo from PD and MSA cases, from recombinant α-synuclein seeded with PD, DLB, or MSA tissue, or from amyloids formed in vitro (Cullinane et al., 2025; Lövestam et al., 2021; Schweighauser et al., 2020; Yang et al., 2023; Yang et al., 2022), we observed structural conservation of a short segment within the NAC region. Specifically, the peptide (^66^VGGAVVT^72^) adopts a highly conserved conformation across these structurally diverse polymorphs, with a backbone RMSD of 1.2 Å (Figure 1A). This shared structural motif is a potential target for inhibition. Literature supports this observation, as a 12-residue segment, ⁷¹VTGVTAVAQKTV⁸², has been reported to be necessary and sufficient for fibril formation (Giasson et al., 2001), while deletion of the shorter ⁶⁶VGGAVVTGV⁷⁴ segment abolishes aggregation and cytotoxicity, establishing this region as a minimal aggregation-critical core (Du et al., 2003). This region also emerged as a critical segment in a study of the aggregation propensity of peptides that tile the α-synuclein sequence (El-Agnaf et al., 2004). Structural studies of the NACore fragment, ⁶⁸GAVVTGVTAVA⁷⁸, revealed that this segment forms steric zipper β-sheet within the fibril spine (Rodriguez et al., 2015). This observation is further supported by near-atomic structures of full-length α-synuclein fibrils, in which NAC and NACore-adjacent residues form the fibril core (Eisenberg and Sawaya, 2017; Guerrero-Ferreira et al., 2019; Li et al., 2018). Short NAC-derived peptides have accordingly been developed both as aggregation modulators and as minimal models for probing sequence-specific interactions (Allen et al., 2023; Bodles et al., 2004; Duan et al., 2025; Sangwan et al., 2020). Modified peptides spanning the α-synuclein binding region (residues 69–72) were shown to bind full-length α-synuclein and block its assembly into both early oligomers and mature amyloid-like fibrils (El-Agnaf et al., 1998). Conversely, GAV-motif peptides from within residues 66–74 accelerate fibrillization of wild-type α-synuclein and induce it in charge-incorporated mutants, but have no effect on a GAV-deficient mutant, demonstrating that the interaction is strictly sequence-specific (Du et al., 2006). Simulations of an amphiphilic NAC-derived peptide further indicate that hydrophilic neighboring groups are required to recruit enough co-solvent to weaken the hydrophobic contacts and disrupt the β-sheet, whereas a purely hydrophobic analogue resists disruption (Galamba, 2022). Based on the centrality of this segment to α-synuclein misfolding, we hypothesized that it could be targeted for modulating α-synuclein aggregation.

**Figure 1.**
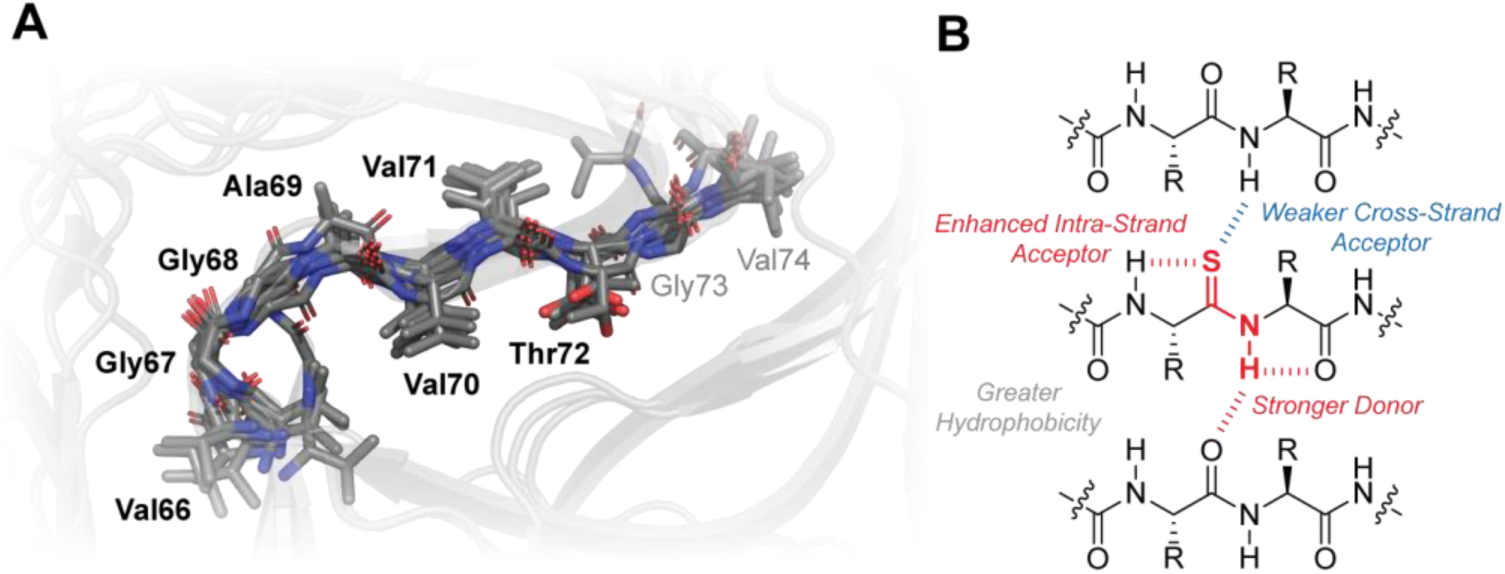
(**A**) (^66^VGGAVVTGV^74^) amino acid sequence alignment within representative α-synuclein fibril structures (PDB: 9cx6, 8a9l, 8cz6, 9euu, 7ozh, 8uka, 8h03, 8pix). (**B**) Effects of thioamide substitution on peptide hydrogen-bonding networks in the context of parallel β-sheets, such as those in amyloid fibers.

Peptides offer great promise in addressing protein aggregation, thanks in part to their size, functional group density, and modularity, which together allow them target critical surfaces and interfaces with specificity that can be difficult to achieve with smaller molecules (Arkin et al., 2014; Cao et al., 2022; Murray et al., 2022a; Nazzaro et al., 2023; Patgiri et al., 2008). For example, macrocyclic β-sheet mimics can lock amyloidogenic strands into a β-hairpin or β-arch, preorganizing them for β-sheet association and lowering the entropic penalty of docking onto a growing cross-β edge, which improves potency over unconstrained sequences (Lu et al., 2019) and improves stability (Freire and Gellman, 2009). Macrocyclic peptides have been developed to target various misfolded proteins, including Aβ, tau, α-synuclein, IAPP, and prion proteins (Cheng et al., 2012). Such molecules can be further optimized by blocking one hydrogen-bonding edge, which can cap amyloid assemblies (Rajewski et al., 2023; Zheng et al., 2011). These strategies show promise for inhibiting α-synuclein assembly (Kritzer et al., 2009; Spanopoulou et al., 2018; Hornung et al., 2025).

Optimized sequence/structural motifs are frequently combined with a range of backbone modifications to disrupt the key hydrogen-bonding interactions that stabilize protein aggregates, including amyloid fibers. For example, amide-to-ester substitutions can reduce β-sheet assembly (Truex et al., 2016) by attenuating backbone hydrogen bonds (Deechongkit et al., 2004a; Deechongkit et al., 2004b; Zheng and Newberry, 2026). Amidines, which substitute the carbonyl oxygen with nitrogen, are tolerated as hydrogen-bond donors but abolish the edge-to-edge hydrogen bonding required for fibril growth (O’Brien et al., 2025). *N*-Amination, which a backbone amide N–H with an N–NH₂ group, stabilizes an extended, sheet-competent conformation but removes a cross-strand donor on one edge, disrupting fibril assembly (Makwana et al., 2021). Among these approaches, *N*-methylation is perhaps the most common modification, which blocks inter-strand hydrogen bonding and inhibits fibril formation (Cao et al., 2020; Chatterjee et al., 2008; Kokkoni et al., 2006; Madine et al., 2008; Samdin et al., 2024; Samdin et al., 2025; Yan et al., 2013).

We propose that thioamides can also serve as useful amide bond isosteres for optimizing peptide modulators of protein aggregation (Figure 1B). Thioamides have attracted significant attention as amide bond isosteres because they largely preserve the resonance stabilization, planarity, and overall hydrogen-bonding framework of native amide bonds (Choudhary and Raines, 2011; Fiore et al., 2022; Fiore et al., 2021; Ghosh et al., 2023; Mahanta et al., 2019; Petersson et al., 2014; Phan et al., 2026; Walters et al., 2017). However, the distinct electronic structure of thioamides perturbs their hydrogen-bonding interactions in ways that are uniquely well-suited to targeting misfolded proteins. For example, thioamides are stronger hydrogen-bond donors due to the lower p*K_a_* of the thioamide N–H relative to oxoamides (Bordwell, 1988), which we hypothesize could be exploited to increase the affinity of a peptide for interacting with the exposed edge of a misfolded β-strand. In contrast, thioamides are generally weaker hydrogen-bond acceptors because they bear less negative charge on sulfur (Choudhary and Raines, 2011; Min et al., 1998), which can destabilize protein interfaces in which the thioamide acts as a hydrogen-bond acceptor (Culik et al., 2012; Walters et al., 2017). We hypothesize that this reduced capacity to accept hydrogen bonds could be exploited to disrupt protein aggregation by disfavoring the recruitment of additional monomers to a growing self-assembly, such as the exposed end of an amyloid fibril.

Intriguingly, recent studies show that thioamides can actually be stronger hydrogen-bond acceptors than oxoamides in cases where the hydrogen-bond donor approaches perpendicularly to the (thio)carbonyl bond axis (Lampkin and VanVeller, 2021; Zheng and Newberry, 2025); this arrangement is found in C5 hydrogen bonds, which occur frequently within individual protein β-strands (Bartlett and Woolfson, 2016; Newberry and Raines, 2016; Zheng and Newberry, 2026), including those in misfolded proteins (Gallagher-Jones et al., 2018; Murray et al., 2022b). The distinct effects of thioamide incorporation on hydrogen bonds received from different directions results in part from the distinct electronic distribution around the sulfur atom in thioamides (Lampkin and VanVeller, 2021; Newberry et al., 2013). We hypothesize that by strengthening intra-strand C5 hydrogen bonds, thioamides could increase the β-strand propensity of a linear peptide segment, thereby preorganizing the molecule to interact with misfolded proteins. Thioamides therefore have the potential to increase engagement with misfolded proteins, both through secondary structure preorganization and enhanced hydrogen-bond donation. Once bound to the misfolded species, attenuated cross-strand interactions could reduce further self-assembly. However, thioamide substitution can also perturb amide bond polarizability and affect overall molecular polarity (Byerly-Duke and VanVeller, 2024; Fiore et al., 2021; Ghosh et al., 2023; Newberry et al., 2013; Yanagawa et al., 2026), which could confound the predicted effects of thioamide substitution on key hydrogen-bonding patterns. We therefore sought to determine how thioamide incorporation would affect the ability of short peptides to engage with misfolded proteins like α-synuclein.

## Results & Discussion

In this study, we targeted a structurally conserved motif within α-synuclein fibers using backbone-modified peptides to test the effects of thioamide incorporation on protein–protein interactions in α-synuclein misfolding. Specifically, the ^66^VGGAVVT^72^ peptide within the NAC region adopts similar conformations across a wide variety of misfolded conformational strains (Figure 1A) (Cullinane et al., 2025; Lövestam et al., 2021; Schweighauser et al., 2020; Yang et al., 2023; Yang et al., 2022), suggesting that it plays a key role in α-synuclein misfolding. Indeed, prior deletion studies have shown that ^66^VGGAVVTGV^74^ plays a key role in both fibrillization and cytotoxicity (Du et al., 2003; Galamba, 2022). Moreover, GAVVT-containing peptide can accelerate fibrillization of wild-type α-synuclein, whereas incorporating charged residues within the motif instead inhibits assembly (Du et al., 2006). However, identifying appropriate substitutions that convert a pro-aggregation peptide into an inhibitor requires empirical optimization. In contrast, backbone modification offers an opportunity to control the activity of these peptides without having to optimize the sequence. We therefore sought to exploit backbone modification, specifically thioamide substitution, to control the activity of the heptapeptide ^66^VGGAVVT^72^, a minimal template that contains the core GAVVT motif in which the conserved fibril core motif is preserved.^13^ The length of the sequence also facilitates synthetic incorporation of backbone modifications, which can present synthetic challenges (Newberry et al., 2015; Walters et al., 2017).

We first compared the aggregation kinetics of this peptide (Pep1-O; Figure 2A) to an analog with a single thioamide substitution between Val71 and Thr72 (Pep1-S1; Figure 2A). We studied peptide aggregation using the turbidity of the solution over time, since peptides this length generally lack the large, rigid hydrophobic surfaces necessary for detection by fluorogenic dyes such as thioflavin T (Du et al., 2006; Groenning, 2010). Under our experimental conditions, Pep1-O aggregates slowly, gradually increasing in turbidity over approx. 20 h (Figure 2B). The result is consistent with what has been reported previously (Du et al., 2006; El-Agnaf et al., 1998). In comparison, Pep1-S1 aggregates more rapidly, reaching a plateau in turbidity within approx. 4 hours. Thioamide substitution therefore appears sufficient to drive this peptide toward the aggregated state, possibly though a confluence of enhanced intra- and cross-strand hydrogen bonding, as well as peptide solvophobicity (Fiore et al., 2022; Newberry et al., 2015; Walters et al., 2017; Zheng and Newberry, 2025). To further test the effects of thioamide substitution on the propensity of the short peptide to aggregate, we mixed equal amounts of the modified (Pep1-S1) and unmodified and found that aggregation kinetics were intermediate between pure samples of either peptide (Figure 2B).

**Figure 2.**
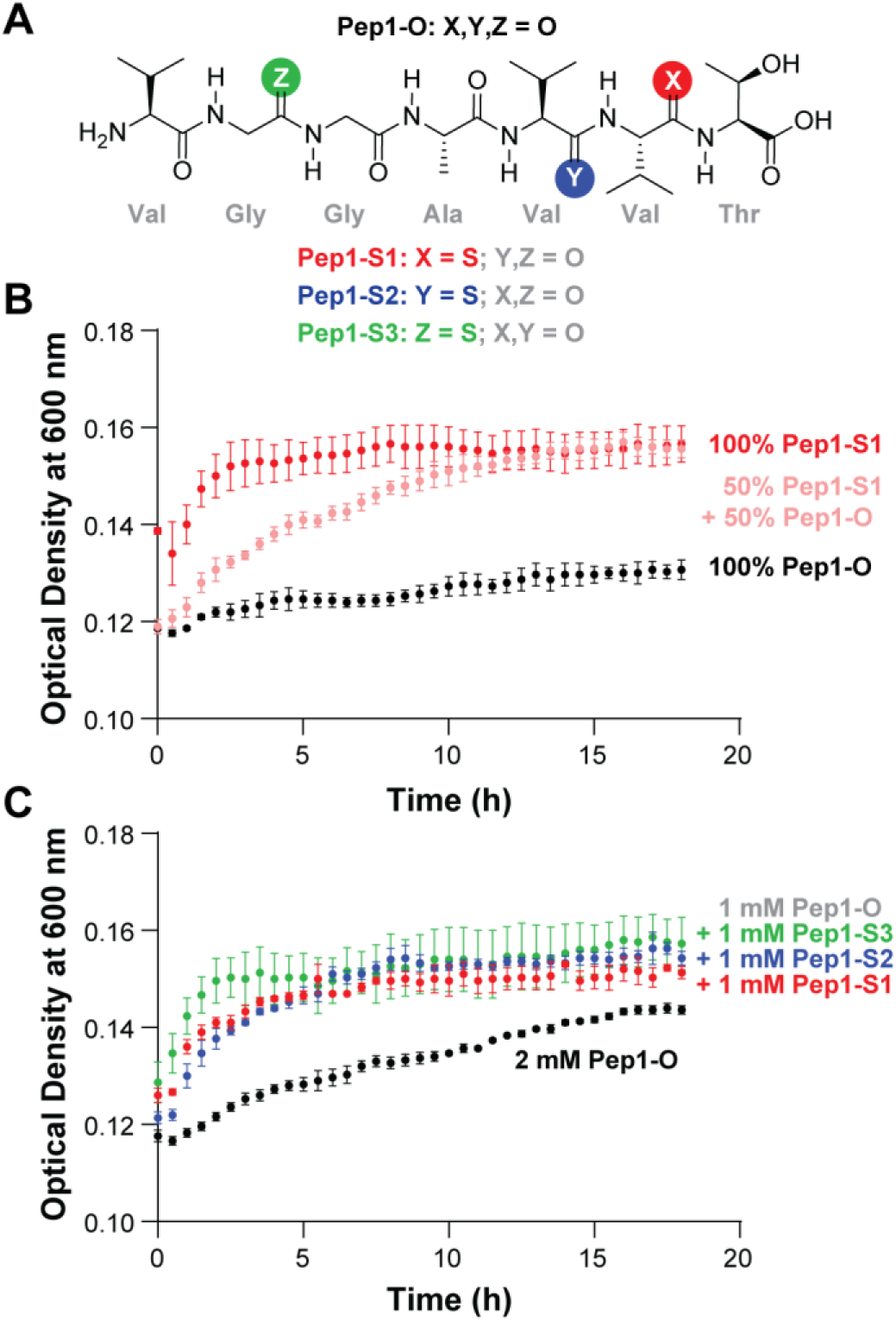
(**A**) NAC-derived peptides and the chemical modifications introduced to study their effect on fibril formation. (**B**) Aggregation kinetics for mixtures of Pep1-O and Pep1-S1. (**C**) Aggregation kinetics of NAC-derived peptides, Pep1-S1, Pep1-S2, and Pep1-S3.

We next asked if the effect of thioamide substitution was specific to the linkage between Val71 and Thr72. To test this, we generated a series of three isomeric thiopeptides (Pep1-S1, Pep1-S2, and Pep1-S3; Figure 2A), in which thioamides were placed at three different backbone positions. The aggregation kinetics of 1:1 mixtures between each thiopeptide and the unmodified parent (Pep1-O) were indistinguishable from one another (Figure 2C). Thioamide substitution therefore likely promotes peptide aggregation by strengthening common molecular driving forces (e.g., hydrogen bonding, hydrophobicity) across different substitution sites within the sequence. These results indicate that thioamide substitution is sufficient to increase the peptide’s population of the aggregated state.

Given the ability of thioamide substitution to promote the aggregated state of a short peptide like Pep1-O, we next asked whether this substitution would similarly promote engagement of the peptide with the aggregated state of full-length α-synuclein. To test this hypothesis, we incubated a fixed concentration of full-length α-synuclein with increasing concentrations of either Pep1-O or Pep1-S1 (Figure 3A) and recorded the rate of aggregation via thioflavin T fluorescence (Figure 3B/C). Full-length α-synuclein forms ThT-positive aggregates within approx. 5 days under our experimental conditions. The addition of either Pep1-O or Pep1-S1 was sufficient to accelerate aggregation in a dose-dependent manner (Figure 3B/C), even though the peptide is only 5% the size of full-length α-synuclein, suggesting the importance of this segment. Because the addition of peptide shortens the lag time before aggregates are detected, we speculate that the peptides facilitate early assembly steps, potentially through local concentration effects, transient interactions with full-length monomers, or surface-mediated conformational conversion (Buell et al., 2014; Vadukul et al., 2023). Consistent with the effects of thioamide substitution on the aggregation of the peptide by itself, Pep1-S1 was significantly more effective at accelerating α-synuclein aggregation than its unmodified counterpart, Pep1-O (Figure 3D), despite the fact that these peptides differ in only a single atom. The combined effects of thioamide substitution on backbone hydrogen bonding and hydrophobicity (Ghosh et al., 2023; Ono et al., 2012) are therefore sufficient to enhance association of this short peptide with α-synuclein aggregates.

**Figure 3.**
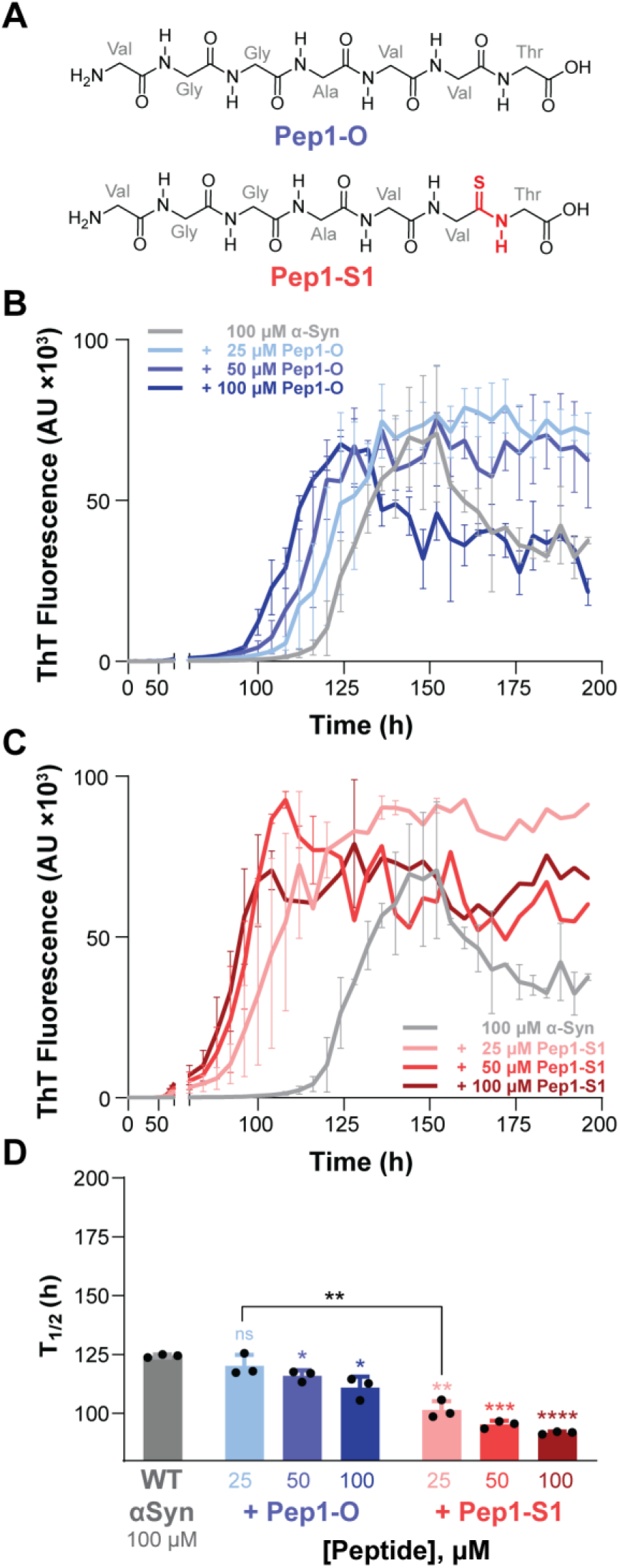
(**A**) Chemical structures of Pep1-O and Pep1-S. (**B**,**C**) α-Synuclein aggregation kinetics in the presence of Pep1-O (**B**) or Pep1-S (**C**), monitored by ThT fluorescence over time. (**D**) Change in aggregation kinetics of full-length α-synuclein in the presence of NAC-derived peptides, measured by the time required to achieve half maximal ThT fluorescence (T_1/2_). Data are presented with triplicate as mean ± SEM, and statistical significance is relative to the no-peptide control, except where indicated. * = p < 0.05, ** = p < 0.01, *** = p < 0.001, **** = p < 0.0001, n.s. = not significant

Encouraged by the ability of thioamides to enhance association with the aggregated state of α-synuclein, we next asked if we could exploit thioamides to improve an aggregation inhibitor. Because the NAC-derived peptide is amyloidogenic in isolation and adopts a conserved β-strand conformation within ex vivo α-synuclein fibrils, we reasoned that targeted backbone modifications within this segment could perturb disease-relevant β-sheet packing. However, thioamide substitution alone enhanced the aggregation propensity of the NAC peptide. This accelerating effect may arise from a shift in the balance of backbone hydrogen-bonding interactions, as thioamides can strengthen hydrogen-bond donation while weakening hydrogen-bond acceptance. We therefore asked whether adding a second backbone modification could sufficiently increase the local perturbation of β-sheet geometry and convert this aggregation-promoting effect into an inhibitory strategy. Hence, we paired thioamide substitution with *N*-methylation, a synthetically accessible modification that removes a backbone hydrogen-bond donor and introduces steric bulk (Chatterjee et al., 2013).

We reasoned that *N*-methylation would most effectively complement thioamide substitution when incorporated onto the same edge of the β-sheet as the thioamide sulfur (Figure 4A); this would position the two hydrogen-bond disrupting groups on the same edge of the β-sheet, while preserving the ability of the unmodified edge to engage full-length α-synuclein and capitalizing on the ability of thioamides to enhance that interaction. We therefore introduced an *N*-methyl group at Gly67 (Figure 4A). As expected, addition of this *N*-methylated peptide (Pep2-O) to full-length α-synuclein was sufficient to slow aggregation (Figure 4B/D). Remarkably, the *N*-methylated thiopeptide (Pep2-S) substantially delayed the aggregation of full-length α-synuclein, even at substoichiometric concentrations (Figure 4C/D). For comparison, short peptide inhibitors of protein aggregation generally require molar excess over the target protein in order to slow aggregation (Cao et al., 2020; Lu et al., 2019; Rajewski et al., 2023). The delay in aggregation observed at 0.25 molar equivalents suggests that thioamide substitution is potent at disrupting cross-β amyloid and can be used as a design element for peptide-based inhibitor design. Furthermore, electron microscopy indicates that fibrils formed in the presence of either Pep2-O or Pep2-S are indistinguishable from those formed by α-synuclein alone (Figure S7), suggesting that these inhibitors primarily modulate aggregation kinetics without measurably altering the final fibril morphology under the conditions tested.

**Figure 4.**
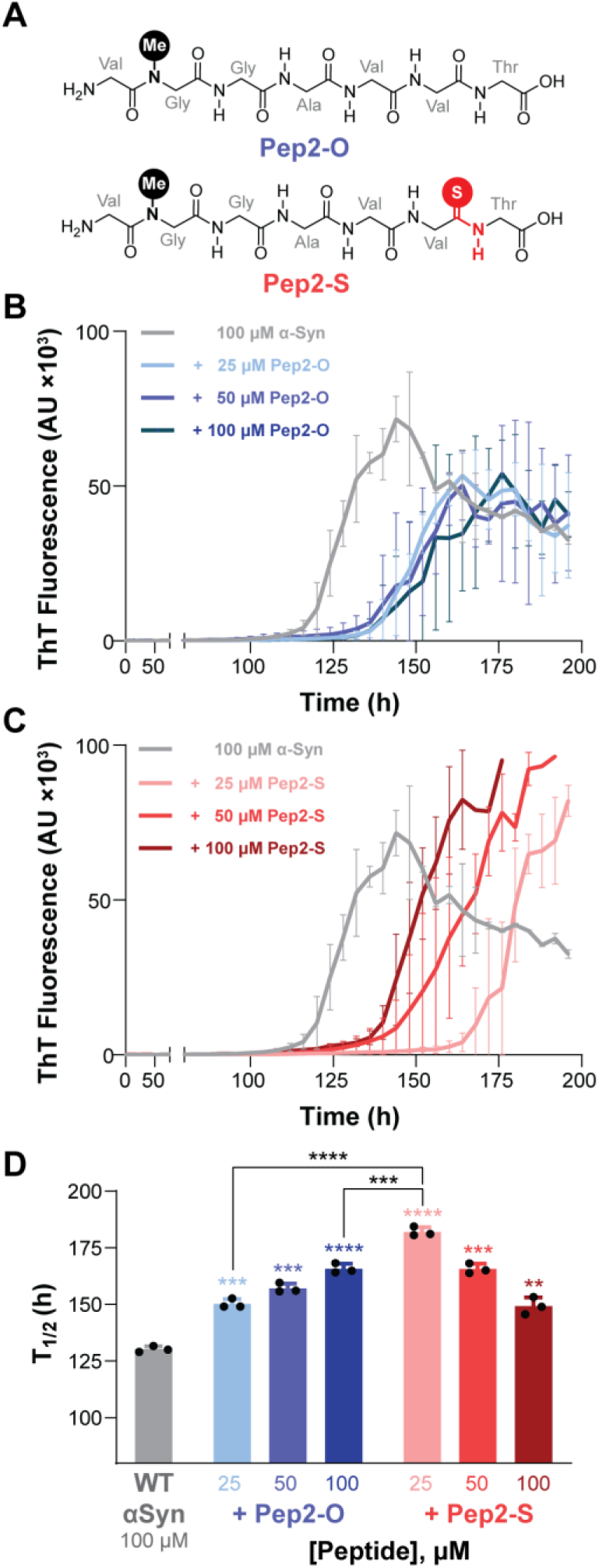
(**A**) Chemical structures of Pep2-O and Pep2-S. (**B**, **C**) α-Synuclein aggregation kinetics in the presence of either Pep2-O (**B**) or Pep2-S (**C**), as measured by changes in ThT fluorescence over time. (**D**) Change in aggregation kinetics of full-length α-synuclein in the presence of NAC-derived peptides, measured by the time required to achieve half maximal ThT fluorescence (T_1/2_). Data are presented with triplicate as mean ± SEM, and statistical significance is relative to the no-peptide control, except where indicated. * = p < 0.05, ** = p < 0.01, *** = p < 0.001, **** = p < 0.0001, n.s. = not significant

Interestingly, the inhibitory activity of Pep2-S decreases with increasing concentration (Figure 4C/D), contrary to the expected dose dependence. Because thioamide substitution reduces local hydration and increases polarizability (Ghosh et al., 2023), we speculate that Pep2-S may exhibit increased concentration-dependent self-association relative to Pep2-O. This could reduce the concentration of freely available inhibitor or alter its interactions with α-synuclein, potentially accounting for the diminished inhibitory activity observed at higher Pep2-S concentrations.

To probe the synergy of *N*-methylation and backbone thioamide substitution, we generated a second inhibitor pair in which the *N*-methyl group is incorporated at Gly68, which would place the methyl group on the opposite edge of the β-sheet from the thioamide sulfur (Pep3-O and Pep3-S; Figure 5A). We hypothesize that this arrangement should neutralize the effects of both modifications, since the presence of disrupting groups on both edges of the β-sheet are expected to hinder association with full-length α-synuclein. Consistent with *N*-methylation at Gly67 (Pep2-O), *N*-methylation of Gly68 (Pep3-O) was sufficient to convert the VGGAVVT peptide into an inhibitor of α-synuclein aggregation (Figure 5B/D). Pep3-O was quantitatively less effective as an inhibitor relative to Pep2-O (Figure 4B/D), suggesting position-specific effects of backbone methylation. Importantly, Pep3-S had minimal effect on the rate of α-synuclein aggregation (Figure 5C/D), consistent with our hypothesis that *N*-methylation and backbone thioamide substitution must be positioned on the same edge of the β-sheet to act synergistically in disrupting the cross-β hydrogen bonding network in order to inhibit protein aggregation.

**Figure 5.**
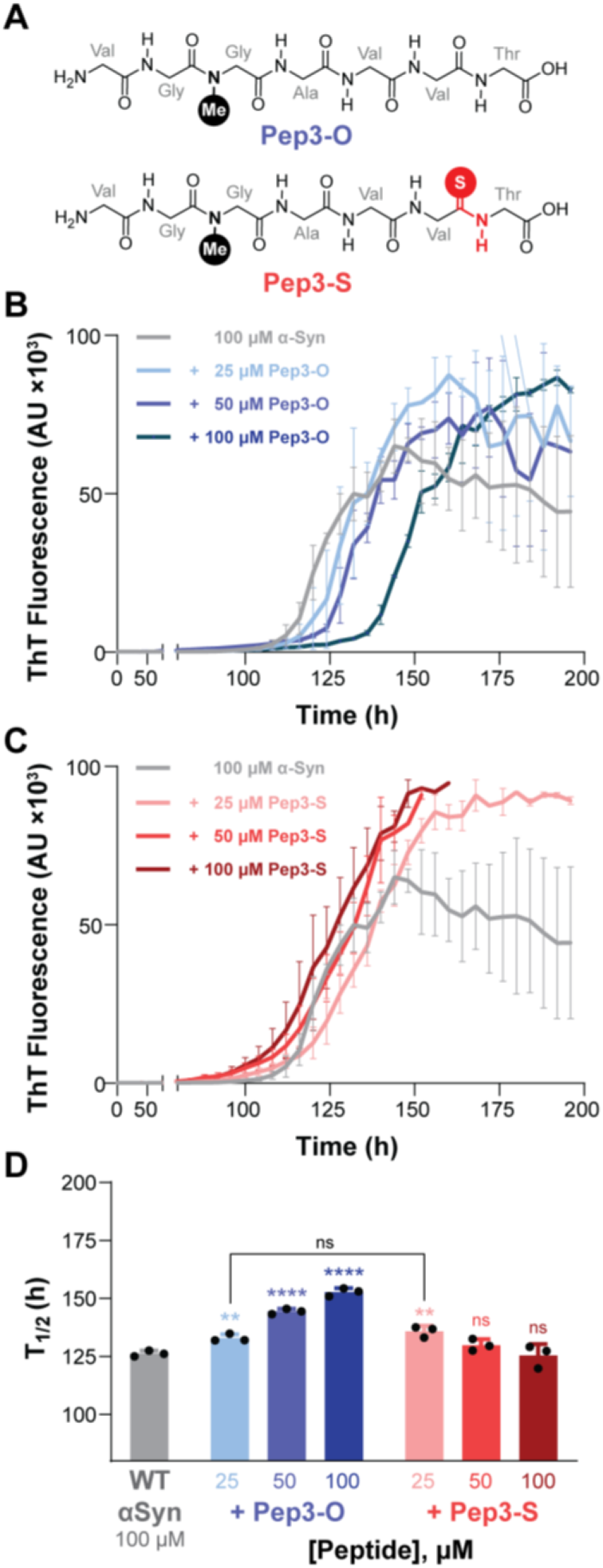
(**A**) Chemical structures of Pep3-O and Pep3-S. (**B**, **C**) α-Synuclein aggregation kinetics in the presence of either Pep3-O (**B**) or Pep3-S (**C**), as measured by changes in ThT fluorescence over time. (**D**) Change in aggregation kinetics of full-length α-synuclein in the presence of NAC-derived peptides, measured by the time required to achieve half maximal ThT fluorescence (T_1/2_). Data are presented with triplicate as mean ± SEM, and statistical significance is relative to the no-peptide control, except where indicated. * = p < 0.05, ** = p < 0.01, *** = p < 0.001, **** = p < 0.0001, n.s. = not significant

Intrigued by the anti-aggregation activity of Pep2-O/S in vitro, we next asked whether these peptides could also reduce seeded aggregation of α-synuclein in cultured cells. When aggregated α-synuclein is introduced to cells expressing α-synuclein, the exogenous aggregates can seed the aggregation of the endogenous protein into subcellular puncta, which can be detected by imaging if the endogenous protein is either fused to a fluorescent reporter or can be stained for immunofluorescence (Figure 6A). Quantification of the puncta therefore serves as a measurement of the cellular seeding activity of α-synuclein aggregates (Maina et al., 2022; Reis et al., 2024; Sang et al., 2021; Woerman et al., 2015; Yamasaki et al., 2019).

**Figure 6.**
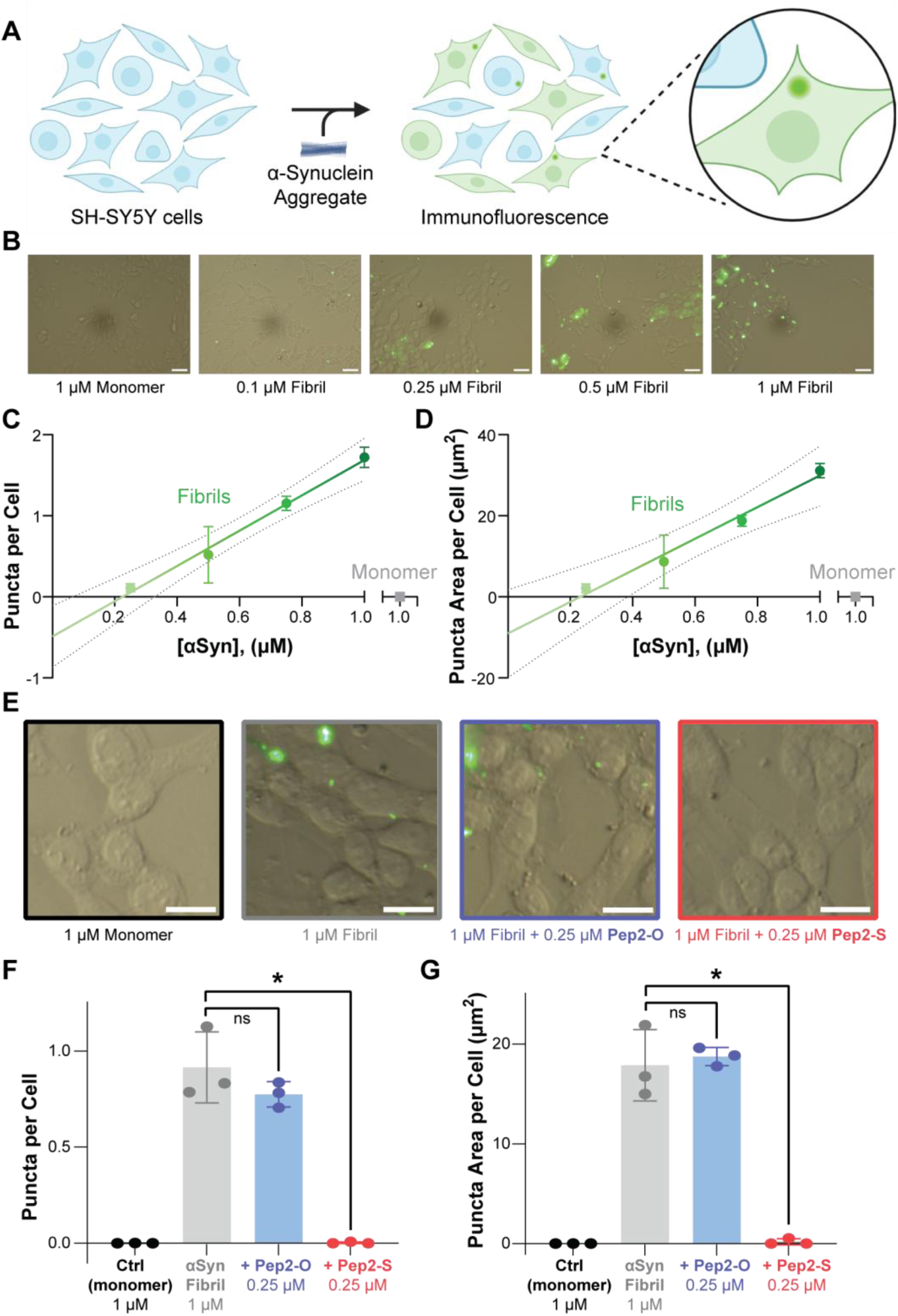
Cellular aggregation of α-synuclein seeded in the presence or absence of NAC-derived peptide inhibitors. (**A**) Overview of the assay for seeded α-synuclein aggregation in SH-SY5Y cells. (**B**) Representative merged brightfield and fluorescence micrographs of SH-SY5Y cells treated with varying concentrations of α-synuclein fibrils. Scale bars are 30 μm. (**C-D**) Quantification of the aggregation observed during the experiment in **B**. (**E**) Representative merged brightfield and fluorescence micrographs of SH-SY5Y cells treated with α-synuclein fibrils formed in the presence of NAC-derived peptides. Scale bars are 10 μm. (**F-G**) Quantification of the puncta observed during the experiment in **E**. Data are collected triplicate, and statistical significance is indicated as *p < 0.05, and n.s. (not significant).

To validate this approach, we introduced varying concentrations of preformed α-synuclein aggregates into SH-SY5Y neuroblastoma cells (Choi et al., 2020; Furthmann et al., 2023; Woerman et al., 2015; Xicoy et al., 2017) and quantified the number and size of intracellular puncta after 6 days (Sang et al., 2021). Within a concentration range of 0.25–1.0 μM exogenous α-synuclein aggregate, the cells demonstrated a linear increase in the quantity of intracellular puncta (Figure 6B-D). In contrast, addition of 1 μM α-synuclein monomer did not change the amount of puncta relative to untreated controls (Figure 6B-D), highlighting the sensitivity of this assay to the seeding activity of the exogenous aggregates.

We then compared the seeding activity of α-synuclein aggregates that had been formed in the presence of 0.25 eq. of either Pep2-O or Pep2-S. Consistent with its greater inhibitory effect in vitro, Pep2-S was significantly more effective at reducing the cellular seeding activity of aggregated α-synuclein than was Pep2-O (Figure 6E-G). These results further emphasize the capacity for synergistic thioamide substitution and *N*-methylation to improve peptide inhibitors of protein aggregation. Thioamide substitution therefore presents a complementary strategy to other backbone modifications that control protein assembly, such as *N*-amination, which has been used to convert amyloidogenic tau peptides into soluble β-strand mimics that similarly block cellular seeding (Makwana et al., 2021). However, the concentration dependence observed here (Figure 4) emphasizes that these peptides must be used within an appropriate inhibitory concentration window, as higher concentrations may favor self-association and reduce inhibitory activity (Griner et al., 2019).

In summary, we demonstrate that minimal amyloidogenic sequences can be rationally reprogrammed through backbone engineering to exert bidirectional control over α-synuclein aggregation. Using a defined NAC-derived motif, ^66^VGGAVVT^72^, thioamide substitution enhances intermolecular interactions and accelerates nucleation, whereas *N*-methylation disrupts β-sheet hydrogen-bonding networks to inhibit fibrillization. Strikingly, their integration yields hybrid peptides that achieve potent, geometry-dependent inhibition of aggregation at substoichiometric concentrations, which can be observed both in vitro and in cellular models. These results highlight an opportunity to target a key segment of α-synuclein in anti-aggregation efforts. More broadly, this study highlights new backbone-engineering approaches to target other amyloidogenic systems and protein–protein interactions, with potential implications for the development of therapeutics for neurodegenerative diseases.

## Acknowledgments

This work was supported by grants from the NIH (R00-NS116679 and R35-GM160477) and the Welch Foundation (F-2116 and F-2298) to RWN. We thank Daeun Noh for assistance with protein expression and Fengyi Gu for assistance with the electron microscopy.

## References

Allen SG, Meade RM, White Stenner LL, Mason JM. Peptide-based approaches to directly target alpha-synuclein in Parkinson’s disease. Mol Neurodegener. 2023; 18(1): 80. 10.1186/s13024-023-00675-8.

Arkin MR, Tang Y, Wells JA. Small-Molecule Inhibitors of Protein-Protein Interactions: Progressing toward the Reality. Chem Biol. 2014; 21(9): 1102–1114. 10.1016/j.chembiol.2014.09.001.

Bartlett GJ, Woolfson DN. On the satisfaction of backbone-carbonyl lone pairs of electrons in protein structures. Protein Sci. 2016; 25(4): 887–897. 10.1002/pro.2896.

Bodles AM, El-Agnaf OMA, Greer B, Guthrie DJS, Irvine GB. Inhibition of fibril formation and toxicity of a fragment of α-synuclein by an N-methylated peptide analogue. Neurosci Lett. 2004; 359(1–2): 89–93. 10.1016/j.neulet.2003.12.077.

Bordwell FG. Equilibrium acidities in dimethyl sulfoxide solution. Acc Chem Res. 1988; 21(12): 456–463. 10.1021/ar00156a004.

Buell AK, Galvagnion C, Gaspar R, Sparr E, Vendruscolo M, Knowles TPJ, Linse S, Dobson CM. Solution conditions determine the relative importance of nucleation and growth processes in α-synuclein aggregation. Proc Natl Acad Sci U S A. 2014; 111(21): 7671–7676. 10.1073/pnas.1315346111.

Byerly-Duke J, VanVeller B. Thioimidate Solutions to Thioamide Problems during Thionopeptide Deprotection. Org Lett. 2024; 26(7): 1452–1457. 10.1021/acs.orglett.4c00035.

Cao L, Coventry B, Goreshnik I, Huang B, Sheffler W, Park JS, Jude KM, Marković I, Kadam RU, Verschueren KHG, Verstraete K, Walsh STR, Bennett N, Phal A, Yang A, Kozodoy L, DeWitt M, Picton L, Miller L, Strauch EM, DeBouver ND, Pires A, Bera AK, Halabiya S, Hammerson B, Yang W, Bernard S, Stewart L, Wilson IA, Ruohola-Baker H, Schlessinger J, Lee S, Savvides SN, Garcia KC, Baker D. Design of protein-binding proteins from the target structure alone. Nature. 2022; 605(7910): 551–560. 10.1038/s41586-022-04654-9.

Cao Q, Boyer DR, Sawaya MR, Ge P, Eisenberg DS. Cryo-EM structure and inhibitor design of human IAPP (amylin) fibrils. Nat Struct Mol Biol. 2020; 27(7): 653–659. 10.1038/s41594-020-0435-3.

Chatterjee J, Gilon C, Hoffman A, Kessler H. N-Methylation of Peptides: A New Perspective in Medicinal Chemistry. Acc Chem Res. 2008; 41(10): 1331–1342. 10.1021/ar8000603.

Chatterjee J, Rechenmacher F, Kessler H. N-Methylation of Peptides and Proteins: An Important Element for Modulating Biological Functions. Angew Chem Int Ed Engl. 2013; 52(1): 254–269. 10.1002/anie.201205674.

Cheng PN, Liu C, Zhao M, Eisenberg D, Nowick JS. Amyloid β-sheet mimics that antagonize protein aggregation and reduce amyloid toxicity. Nat Chem. 2012; 4(11): 927–933. 10.1038/nchem.1433.

Chiti F, Dobson CM. Protein Misfolding, Amyloid Formation, and Human Disease: A Summary of Progress Over the Last Decade. Annu Rev Biochem. 2017; 86(1): 27–68. 10.1146/annurev-biochem-061516-045115.

Choi YR, Kim JB, Kang SJ, Noh HR, Jou I, Joe EH, Park SM. The dual role of c-src in cell-to-cell transmission of α-synuclein. EMBO Rep. 2020; 21(7): e48950. 10.15252/embr.201948950.

Choudhary A, Raines RT. An Evaluation of Peptide-Bond Isosteres. ChemBioChem. 2011; 12(12): 1801–1807. 10.1002/cbic.201100272.

Culik RM, Jo H, DeGrado WF, Gai F. Using Thioamides To Site-Specifically Interrogate the Dynamics of Hydrogen Bond Formation in β-Sheet Folding. J Am Chem Soc. 2012; 134(19): 8026–8029. 10.1021/ja301681v.

Cullinane PW, Yang Y, Chelban V, Goh YY, Ebanks K, Curless T, Wrigley S, de Pablo-Fernández E, Holton JL, Peak-Chew S, Franco C, Woerman AL, Houlden H, Warner TT, Scheres SHW, Goedert M, Jaunmuktane Z. Identical Seeding Characteristics and Cryo-EM Filament Structures in FTLD-Synuclein and Typical Multiple System Atrophy. Neuropathol Appl Neurobiol. 2025; 51(2): e70013. 10.1111/nan.70013.

Deechongkit S, Dawson PE, Kelly JW. Toward Assessing the Position-Dependent Contributions of Backbone Hydrogen Bonding to β-Sheet Folding Thermodynamics Employing Amide-to-Ester Perturbations. J Am Chem Soc. 2004; 126(51): 16762–16771. 10.1021/ja045934s.

Deechongkit S, Nguyen H, Powers ET, Dawson PE, Gruebele M, Kelly JW. Context-dependent contributions of backbone hydrogen bonding to β-sheet folding energetics. Nature. 2004; 430(6995): 101–105. 10.1038/nature02611.

Du HN, Li HT, Zhang F, Lin XJ, Shi JH, Shi YH, Ji LN, Hu J, Lin DH, Hu HY. Acceleration of α-synuclein aggregation by homologous peptides. FEBS Lett. 2006; 580(15): 3657–3664. 10.1016/j.febslet.2006.05.050.

Du HN, Tang L, Luo XY, Li HT, Hu J, Zhou JW, Hu HY. A Peptide Motif Consisting of Glycine, Alanine, and Valine Is Required for the Fibrillization and Cytotoxicity of Human α-Synuclein. Biochemistry. 2003; 42(29): 8870–8878. 10.1021/bi034028+.

Duan J, Zhang H, Sun C. AI-Guided Dual Strategy for Peptide Inhibitor Design Targeting Structural Polymorphs of α-Synuclein Fibrils. Cells. 2025; 14(23): 1921. 10.3390/cells14231921.

Eisenberg DS, Sawaya MR. Structural Studies of Amyloid Proteins at the Molecular Level. Annu Rev Biochem. 2017; 86(1): 69–95. 10.1146/annurev-biochem-061516-045104.

El-Agnaf OMA, Jakes R, Curran MD, Middleton D, Ingenito R, Bianchi E, Pessi A, Neill D, Wallace A. Aggregates from mutant and wild-type α-synuclein proteins and NAC peptide induce apoptotic cell death in human neuroblastoma cells by formation of β-sheet and amyloid-like filaments. FEBS Lett. 1998; 440(1): 71–75. 10.1016/S0014-5793(98)01418-5.

El-Agnaf OMA, Paleologou KE, Greer B, Abogrein AM, King JE, Salem SA, Fullwood NJ, Benson FE, Hewitt R, Ford KJ, Martin FI, Harriott P, Cookson MR, Allsop D. A strategy for designing inhibitors of α-synuclein aggregation and toxicity as a novel treatment for Parkinson’s disease and related disorders. FASEB J. 2004; 18(11): 1315–1317. 10.1096/fj.03-1346fje.

Fiore KE, Patist MJ, Giannakoulias S, Huang CH, Verma H, Khatri B, Cheng RP, Chatterjee J, Petersson EJ. Structural impact of thioamide incorporation into a β-hairpin. RSC Chem Biol. 2022; 3(5): 582–591. 10.1039/D1CB00229E.

Fiore KE, Phan HAT, Robkis DM, Walters CR, Petersson EJ. Incorporating thioamides into proteins by native chemical ligation. Methods Enzymol. 2021; 656: 295–339. 10.1016/bs.mie.2021.04.011.

Freire F, Gellman SH. Macrocyclic Design Strategies for Small, Stable Parallel β-Sheet Scaffolds. J Am Chem Soc. 2009; 131(23): 7970–7972. 10.1021/ja902210f.

Furthmann N, Bader V, Angersbach L, Blusch A, Goel S, Sánchez-Vicente A, Krause LJ, Chaban SA, Grover P, Trinkaus VA, van Well EM, Jaugstetter M, Tschulik K, Damgaard RB, Saft C, Ellrichmann G, Gold R, Koch A, Englert B, Westenberger A, Klein C, Jungbluth L, Sachse C, Behrends C, Glatzel M, Hartl FU, Nakamura K, Christine CW, Huang EJ, Tatzelt J, Winklhofer KF. NEMO reshapes the α-Synuclein aggregate interface and acts as an autophagy adapter by co-condensation with p62. Nat Commun. 2023; 14(1): 8368. 10.1038/s41467-023-44033-0.

Galamba N. Aggregation of a Parkinson’s Disease-Related Peptide: When Does Urea Weaken Hydrophobic Interactions? ACS Chem Neurosci. 2022; 13(12): 1769–1781. 10.1021/acschemneuro.2c00169.

Gallagher-Jones M, Glynn C, Boyer DR, Martynowycz MW, Hernandez E, Miao J, Zee CT, Novikova IV, Goldschmidt L, McFarlane HT, Helguera GF, Evans JE, Sawaya MR, Cascio D, Eisenberg DS, Gonen T, Rodriguez JA. Sub-ångström cryo-EM structure of a prion protofibril reveals a polar clasp. Nat Struct Mol Biol. 2018; 25(2): 131–134. 10.1038/s41594-017-0018-0.

Ghosh P, Raj N, Verma H, Patel M, Chakraborti S, Khatri B, Doreswamy CM, Anandakumar SR, Seekallu S, Dinesh MB, Jadhav G, Yadav PN, Chatterjee J. An amide to thioamide substitution improves the permeability and bioavailability of macrocyclic peptides. Nat Commun. 2023; 14(1): 6050. 10.1038/s41467-023-41748-y.

Giasson BI, Murray IVJ, Trojanowski JQ, Lee VMY. A Hydrophobic Stretch of 12 Amino Acid Residues in the Middle of α-Synuclein Is Essential for Filament Assembly. J Biol Chem. 2001; 276(4): 2380–2386. 10.1074/jbc.M008919200.

Goedert M, Spillantini MG, Del Tredici K, Braak H. 100 years of Lewy pathology. Nat Rev Neurol. 2013; 9(1): 13–24. 10.1038/nrneurol.2012.242.

Goldschmidt L, Teng PK, Riek R, Eisenberg D. Identifying the amylome, proteins capable of forming amyloid-like fibrils. Proc Natl Acad Sci U S A. 2010; 107(8): 3487–3492. 10.1073/pnas.0915166107.

Griner SL, Seidler P, Bowler J, Murray KA, Yang TP, Sahay S, Sawaya MR, Cascio D, Rodriguez JA, Philipp S, Sosna J, Glabe CG, Gonen T, Eisenberg DS. Structure-based inhibitors of amyloid beta core suggest a common interface with tau. eLife. 2019; 8: e46924. 10.7554/eLife.46924.

Groenning M. Binding mode of Thioflavin T and other molecular probes in the context of amyloid fibrils-current status. J Chem Biol. 2010; 3(1): 1–18. 10.1007/s12154-009-0027-5.

Guerrero-Ferreira R, Taylor NMI, Arteni AA, Kumari P, Mona D, Ringler P, Britschgi M, Lauer ME, Makky A, Verasdonck J, Riek R, Melki R, Meier BH, Böckmann A, Bousset L, Stahlberg H. Two new polymorphic structures of human full-length alpha-synuclein fibrils solved by cryo-electron microscopy. eLife. 2019; 8: e48907. 10.7554/eLife.48907.

Han H, Weinreb PH, Lansbury PT Jr. The core Alzheimer’s peptide NAC forms amyloid fibrils which seed and are seeded by beta-amyloid: is NAC a common trigger or target in neurodegenerative disease? Chem Biol. 1995; 2(3): 163–169. 10.1016/1074-5521(95)90071-3.

Hornung S, Vogl DP, Naltsas D, Dalla Volta B, Ballmann M, Marcon B, Syed MMK, Wu Y, Spanopoulou A, Feederle R, Heidrich L, Bernhagen J, Koeglsperger T, Höglinger GU, Rammes G, Lashuel HA, Kapurniotu A. Multi-Targeting Macrocyclic Peptides as Nanomolar Inhibitors of Self- and Cross-Seeded Amyloid Self-Assembly of α-Synuclein. Angew Chem Int Ed Engl. 2025; 64(14): e202422834. 10.1002/anie.202422834.

Jakes R, Spillantini MG, Goedert M. Identification of two distinct synucleins from human brain. FEBS Lett. 1994; 345(1): 27–32. 10.1016/0014-5793(94)00395-5.

Kokkoni N, Stott K, Amijee H, Mason JM, Doig AJ. N-Methylated Peptide Inhibitors of β-Amyloid Aggregation and Toxicity. Optimization of the Inhibitor Structure. Biochemistry. 2006; 45(32): 9906–9918. 10.1021/bi060837s.

Kritzer JA, Hamamichi S, McCaffery JM, Santagata S, Naumann TA, Caldwell KA, Caldwell GA, Lindquist S. Rapid selection of cyclic peptides that reduce alpha-synuclein toxicity in yeast and animal models. Nat Chem Biol. 2009; 5(9): 655–663. 10.1038/nchembio.193.

Lampkin BJ, VanVeller B. Hydrogen Bond and Geometry Effects of Thioamide Backbone Modifications. J Org Chem. 2021; 86(24): 18287–18291. 10.1021/acs.joc.1c02373.

Li B, Ge P, Murray KA, Sheth P, Zhang M, Nair G, Sawaya MR, Shin WS, Boyer DR, Ye S, Eisenberg DS, Zhou ZH, Jiang L. Cryo-EM of full-length α-synuclein reveals fibril polymorphs with a common structural kernel. Nat Commun. 2018; 9(1): 3609. 10.1038/s41467-018-05971-2.

Lövestam S, Schweighauser M, Matsubara T, Murayama S, Tomita T, Ando T, Hasegawa K, Yoshida M, Tarutani A, Hasegawa M, Goedert M, Scheres SHW. Seeded assembly in vitro does not replicate the structures of α-synuclein filaments from multiple system atrophy. FEBS Open Bio. 2021; 11(4): 999–1013. 10.1002/2211-5463.13110.

Lu J, Cao Q, Wang C, Zheng J, Luo F, Xie J, Li Y, Ma X, He L, Eisenberg D, Nowick JS, Jiang L, Li D. Structure-based peptide inhibitor design of amyloid-β aggregation. Front Mol Neurosci. 2019; 12: 54. 10.3389/fnmol.2019.00054.

Luk KC, Kehm V, Carroll J, Zhang B, O’Brien P, Trojanowski JQ, Lee VMY. Pathological α-Synuclein Transmission Initiates Parkinson-like Neurodegeneration in Nontransgenic Mice. Science. 2012; 338(6109): 949–953. 10.1126/science.1227157.

Madine J, Doig AJ, Middleton DA. Design of an N-Methylated Peptide Inhibitor of α-Synuclein Aggregation Guided by Solid-State NMR. J Am Chem Soc. 2008; 130(25): 7873–7881. 10.1021/ja075356q.

Mahanta N, Szantai-Kis DM, Petersson EJ, Mitchell DA. Biosynthesis and Chemical Applications of Thioamides. ACS Chem Biol. 2019; 14(2): 142–163. 10.1021/acschembio.8b01022.

Maina KN, Smet-Nocca C, Bitan G. Using FRET-Based Biosensor Cells to Study the Seeding Activity of Tau and α-Synuclein. Methods Mol Biol. 2023; 2551: 125–145. 10.1007/978-1-0716-2597-2_10.

Makwana KM, Sarnowski MP, Miao J, Lin YS, Del Valle JR. N-Amination Converts Amyloidogenic Tau Peptides into Soluble Antagonists of Cellular Seeding. ACS Chem Neurosci. 2021; 12(20): 3928–3938. 10.1021/acschemneuro.1c00528.

Min BK, Lee HJ, Choi YS, Park J, Yoon CJ, Yu JA. A comparative study on the hydrogen bonding ability of amide and thioamide using near IR spectroscopy. J Mol Struct. 1998; 471(1): 283–288. 10.1016/S0022-2860(98)00488-8.

Murray IVJ, Giasson BI, Quinn SM, Koppaka V, Axelsen PH, Ischiropoulos H, Trojanowski JQ, Lee VMY. Role of α-Synuclein Carboxy-Terminus on Fibril Formation in Vitro. Biochemistry. 2003; 42(28): 8530–8540. 10.1021/bi027363r.

Murray KA, Hu CJ, Griner SL, Pan H, Bowler JT, Abskharon R, Rosenberg GM, Cheng X, Seidler PM, Eisenberg DS. De novo designed protein inhibitors of amyloid aggregation and seeding. Proc Natl Acad Sci U S A. 2022; 119(34): e2206240119. 10.1073/pnas.2206240119.

Murray KA, Evans D, Hughes MP, Sawaya MR, Hu CJ, Houk KN, Eisenberg DS. Extended β-Strands Contribute to Reversible Amyloid Formation. ACS Nano. 2022; 16(2): 2154–2163. 10.1021/acsnano.1c08043.

Murray KA, Hu CJ, Pan H, Lu J, Abskharon R, Bowler JT, Rosenberg GM, Williams CK, Elezi G, Balbirnie M, Faull KF, Vinters HV, Seidler PM, Eisenberg DS. Small molecules disaggregate alpha-synuclein and prevent seeding from patient brain-derived fibrils. Proc Natl Acad Sci U S A. 2023; 120(7): e2217835120. 10.1073/pnas.2217835120.

Nazzaro A, Lu B, Sawyer N, Watkins AM, Arora PS. Macrocyclic β-Sheets Stabilized by Hydrogen Bond Surrogates. Angew Chem Int Ed Engl. 2023; 62(41): e202303943. 10.1002/anie.202303943.

Newberry RW, Raines RT. A prevalent intraresidue hydrogen bond stabilizes proteins. Nat Chem Biol. 2016; 12(12): 1084–1088. 10.1038/nchembio.2206.

Newberry RW, VanVeller B, Guzei IA, Raines RT. n→π* Interactions of Amides and Thioamides: Implications for Protein Stability. J Am Chem Soc. 2013; 135(21): 7843–7846. 10.1021/ja4033583.

Newberry RW, VanVeller B, Raines RT. Thioamides in the collagen triple helix. Chem Commun. 2015; 51(47): 9624–9627. 10.1039/C5CC02685G.

O’Brien EA, Abbasi M, Purslow JA, VanVeller B. The ‘ins’ and ‘outs’ of amidines in β-sheet folding and fibril disaggregation. Chem Sci. 2025; 16(36): 16970–16978. 10.1039/D5SC05902J.

Ono K, Takahashi R, Ikeda T, Yamada M. Cross-seeding effects of amyloid β-protein and α-synuclein. J Neurochem. 2012; 122(5): 883–890. 10.1111/j.1471-4159.2012.07847.x.

Patgiri A, Jochim AL, Arora PS. A Hydrogen Bond Surrogate Approach for Stabilization of Short Peptide Sequences in α-Helical Conformation. Acc Chem Res. 2008; 41(10): 1289–1300. 10.1021/ar700264k.

Petersson EJ, Goldberg JM, Wissner RF. On the use of thioamides as fluorescence quenching probes for tracking protein folding and stability. Phys Chem Chem Phys. 2014; 16(15): 6827–6837. 10.1039/C3CP55525A.

Phan HAT, Chang Y, Barrett TM, Fiore KE, Zhang DY, Grove EJ, Petersson EJ. Stabilized thioamide peptide agonists of the neuropeptide Y type 2 receptor for targeted cancer imaging. RSC Chem Biol. 2026; 7(8): 1597–1604. 10.1039/D6CB00042H.

Polymeropoulos MH, Lavedan C, Leroy E, Ide SE, Dehejia A, Dutra A, Pike B, Root H, Rubenstein J, Boyer R, Stenroos ES, Chandrasekharappa S, Athanassiadou A, Papapetropoulos T, Johnson WG, Lazzarini AM, Duvoisin RC, Di Iorio G, Golbe LI, Nussbaum RL. Mutation in the α-Synuclein Gene Identified in Families with Parkinson’s Disease. Science. 1997; 276(5321): 2045–2047. 10.1126/science.276.5321.2045.

Rajewski BH, Makwana KM, Angera IJ, Geremia DK, Zepeda AR, Hallinan GI, Vidal R, Ghetti B, Serrano AL, Del Valle JR. β-Bracelets: Macrocyclic Cross-β Epitope Mimics Based on a Tau Conformational Strain. J Am Chem Soc. 2023; 145(42): 23131–23142. 10.1021/jacs.3c06830.

Reis PM, Holec SAM, Ezeiruaku C, Frost MP, Brown CK, Liu SL, Olson SH, Woerman AL. Structurally targeted mutagenesis identifies key residues supporting α-synuclein misfolding in multiple system atrophy. J Parkinsons Dis. 2024; 14(8): 1543–1558. 10.3233/JPD-240296.

Rodriguez JA, Ivanova MI, Sawaya MR, Cascio D, Reyes FE, Shi D, Sangwan S, Guenther EL, Johnson LM, Zhang M, Jiang L, Arbing MA, Nannenga BL, Hattne J, Whitelegge JP, Brewster AS, Messerschmidt M, Boutet S, Sauter NK, Gonen T, Eisenberg DS. Structure of the toxic core of α-synuclein from invisible crystals. Nature. 2015; 525(7570): 486–490. 10.1038/nature15368.

Samdin TD, Jones CR, Guaglianone G, Kreutzer AG, Freites JA, Wierzbicki M, Nowick JS. A β-Barrel-like Tetramer Formed by a β-Hairpin Derived from Aβ. Chem Sci. 2024; 15(1): 285–297. 10.1039/D3SC05185D.

Samdin TD, Kreutzer AG, Sahrai V, Wierzbicki M, Nowick JS. α-Methylation Enables the X-ray Crystallographic Observation of Oligomeric Assemblies Formed by a β-Hairpin Peptide Derived from Aβ. J Org Chem. 2025; 90(1): 394–400. 10.1021/acs.joc.4c02344.

Sang JC, Hidari E, Meisl G, Ranasinghe RT, Spillantini MG, Klenerman D. Super-resolution imaging reveals α-synuclein seeded aggregation in SH-SY5Y cells. Commun Biol. 2021; 4(1): 613. 10.1038/s42003-021-02126-w.

Sangwan S, Sahay S, Murray KA, Morgan S, Guenther EL, Jiang L, Williams CK, Vinters HV, Goedert M, Eisenberg DS. Inhibition of synucleinopathic seeding by rationally designed inhibitors. eLife. 2020; 9: e46775. 10.7554/eLife.46775.

Schweighauser M, Shi Y, Tarutani A, Kametani F, Murzin AG, Ghetti B, Matsubara T, Tomita T, Ando T, Hasegawa K, Murayama S, Yoshida M, Hasegawa M, Scheres SHW, Goedert M. Structures of α-synuclein filaments from multiple system atrophy. Nature. 2020; 585(7825): 464–469. 10.1038/s41586-020-2317-6.

Spanopoulou A, Heidrich L, Chen HR, Frost C, Hrle D, Malideli E, Hille K, Grammatikopoulos A, Bernhagen J, Zacharias M, Rammes G, Kapurniotu A. Designed Macrocyclic Peptides as Nanomolar Amyloid Inhibitors Based on Minimal Recognition Elements. Angew Chem Int Ed Engl. 2018; 57(44): 14503–14508. 10.1002/anie.201802979.

Spillantini MG, Schmidt ML, Lee VMY, Trojanowski JQ, Jakes R, Goedert M. α-Synuclein in Lewy bodies. Nature. 1997; 388(6645): 839–840. 10.1038/42166.

Truex NL, Wang Y, Nowick JS. Assembly of Peptides Derived from β-Sheet Regions of β-Amyloid. J Am Chem Soc. 2016; 138(42): 13882–13890. 10.1021/jacs.6b06000.

Uéda K, Fukushima H, Masliah E, Xia Y, Iwai A, Yoshimoto M, Otero DA, Kondo J, Ihara Y, Saitoh T. Molecular cloning of cDNA encoding an unrecognized component of amyloid in Alzheimer disease. Proc Natl Acad Sci U S A. 1993; 90(23): 11282–11286. 10.1073/pnas.90.23.11282.

Vadukul DM, Papp M, Thrush RJ, Wang J, Jin Y, Arosio P, Aprile FA. α-Synuclein Aggregation Is Triggered by Oligomeric Amyloid-β42 via Heterogeneous Primary Nucleation. J Am Chem Soc. 2023; 145(33): 18276–18285. 10.1021/jacs.3c03212.

Volpicelli-Daley LA, Luk KC, Patel TP, Tanik SA, Riddle DM, Stieber A, Meaney DF, Trojanowski JQ, Lee VMY. Exogenous α-Synuclein Fibrils Induce Lewy Body Pathology Leading to Synaptic Dysfunction and Neuron Death. Neuron. 2011; 72(1): 57–71. 10.1016/j.neuron.2011.08.033.

Walters CR, Szantai-Kis DM, Zhang Y, Reinert ZE, Horne WS, Chenoweth DM, Petersson EJ. The effects of thioamide backbone substitution on protein stability: a study in α-helical, β-sheet, and polyproline II helical contexts. Chem Sci. 2017; 8(4): 2868–2877. 10.1039/C6SC05580J.

Woerman AL, Stöhr J, Aoyagi A, Rampersaud R, Krejciova Z, Watts JC, Ohyama T, Patel S, Widjaja K, Oehler A, Sanders DW, Diamond MI, Seeley WW, Middleton LT, Gentleman SM, Mordes DA, Südhof TC, Giles K, Prusiner SB. Propagation of prions causing synucleinopathies in cultured cells. Proc Natl Acad Sci U S A. 2015; 112(35): E4949–E4958. 10.1073/pnas.1513426112.

Xicoy H, Wieringa B, Martens GJM. The SH-SY5Y cell line in Parkinson’s disease research: a systematic review. Mol Neurodegener. 2017; 12(1): 10. 10.1186/s13024-017-0149-0.

Yamasaki TR, Holmes BB, Furman JL, Dhavale DD, Su BW, Song ES, Cairns NJ, Kotzbauer PT, Diamond MI. Parkinson’s disease and multiple system atrophy have distinct α-synuclein seed characteristics. J Biol Chem. 2019; 294(3): 1045–1058. 10.1074/jbc.RA118.004471.

Yan LM, Velkova A, Tatarek-Nossol M, Rammes G, Sibaev A, Andreetto E, Kracklauer M, Bakou M, Malideli E, Göke B, Schirra J, Storr M, Kapurniotu A. Selectively N-methylated soluble IAPP mimics as potent IAPP receptor agonists and nanomolar inhibitors of cytotoxic self-assembly of both IAPP and Aβ40. Angew Chem Int Ed Engl. 2013; 52(39): 10378–10383. 10.1002/anie.201302840.

Yanagawa ESK, Fiore KE, Francis DY, Lesneski A, Chang Y, Roose B, Christianson DW, Sato K, Petersson EJ. Improved Protein Semi-Synthesis Enables Biophysical Studies of Thioamide Destabilization of β-Sheet Interactions. bioRxiv. 2026. 10.64898/2026.07.30.741859.

Yang Y, Garringer HJ, Shi Y, Lövestam S, Peak-Chew S, Zhang X, Kotecha A, Bacioglu M, Koto A, Takao M, Spillantini MG, Ghetti B, Vidal R, Goedert M. New SNCA mutation and structures of α-synuclein filaments from juvenile-onset synucleinopathy. Acta Neuropathol. 2023; 145(5): 561–572. 10.1007/s00401-023-02550-8.

Yang Y, Shi Y, Schweighauser M, Zhang X, Kotecha A, Murzin AG, Garringer HJ, Cullinane PW, Saito Y, Foroud T, Warner TT, Hasegawa K, Vidal R, Murayama S, Revesz T, Ghetti B, Hasegawa M, Lashley T, Scheres SHW, Goedert M. Structures of α-synuclein filaments from human brains with Lewy pathology. Nature. 2022; 610(7933): 791–795. 10.1038/s41586-022-05319-3.

Zheng H, Newberry RW. Thioamides in C5 Hydrogen Bonds: Implications for Protein β-Strands. J Org Chem. 2025; 90(39): 13984–13988. 10.1021/acs.joc.5c01310.

Zheng H, Newberry RW. Backbone double-mutant cycle analysis quantifies hydrogen-bond energies in proteins. Protein Sci. 2026; 35(6): e70601. 10.1002/pro.70601.

Zheng J, Liu C, Sawaya MR, Vadla B, Khan S, Woods RJ, Eisenberg D, Goux WJ, Nowick JS. Macrocyclic β-Sheet Peptides That Inhibit the Aggregation of a Tau-Protein-Derived Hexapeptide. J Am Chem Soc. 2011; 133(9): 3144–3157. 10.1021/ja110545h.

