## Supplementary Information for "Backbone Thioamide Substitution Enhances the Activity of Short Peptides in Modulating the Aggregation of α-Synuclein"


| Page | Contents |
| --- | --- |
| S1 | Table of Contents |
| S2 | **Figure S1.** ThT fluorescence kinetics demonstrate that Pep1-O peptide modulate α-synuclein monomer aggregation. |
| S2 | **Figure S2.** ThT fluorescence kinetics demonstrate that Pep1-S1 peptide modulate α-synuclein monomer aggregation. |
| S3 | **Figure S3.** ThT fluorescence kinetics demonstrate that Pep2-O peptide modulate α-synuclein monomer aggregation. |
| S3 | **Figure S4.** ThT fluorescence kinetics demonstrate that Pep2-S peptide modulate α-synuclein monomer aggregation. |
| S4 | **Figure S5.** ThT fluorescence kinetics demonstrate that Pep3-O peptide modulate α-synuclein monomer aggregation. |
| S4 | **Figure S6.** ThT fluorescence kinetics demonstrate that Pep3-S peptide modulate α-synuclein monomer aggregation. |
| S5 | **Figure S7.** Transmission electron microscopy (TEM) of α-synuclein treated with Pep2-O or Pep2-S |
| S6 | General Synthetic Procedures |
| S6 | Peptide Synthetic Procedures |
| S6 | Expression and purification of α-synuclein |
| S7 | Analytical Procedures |
| S8-S10 | HPLC/Analytical LC–MS traces for purified peptides |
| S11 | References |

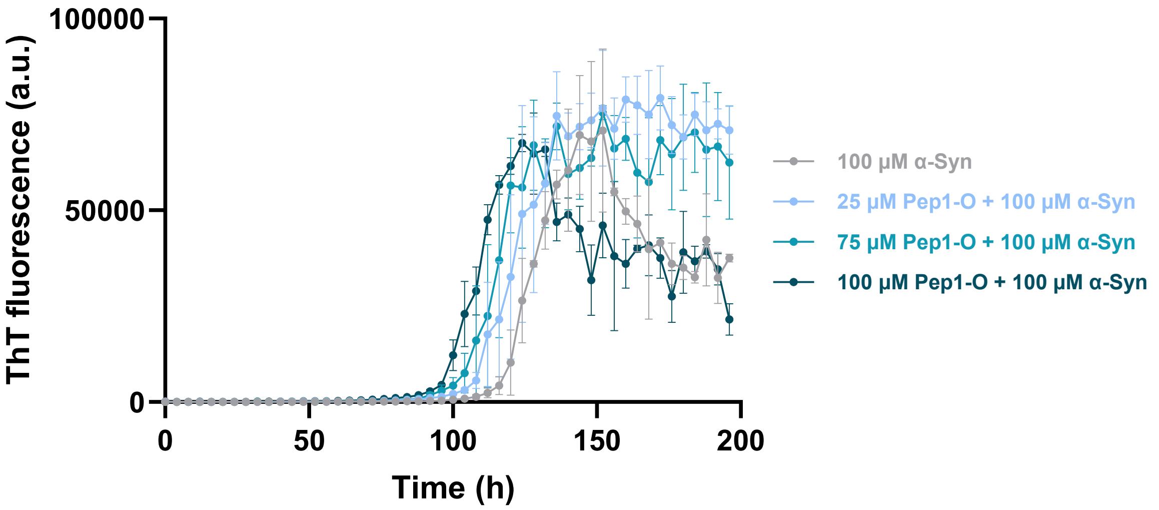


Figure S1. ThT fluorescence kinetics monitoring the aggregation of α-synuclein monomers (100 μM) in the absence or presence of Pep1-O peptide. Direct comparison shows the effects of 25, 75, and 100 μM Pep1-O on α-synuclein aggregation. Measurements were performed in triplicate (n = 3).

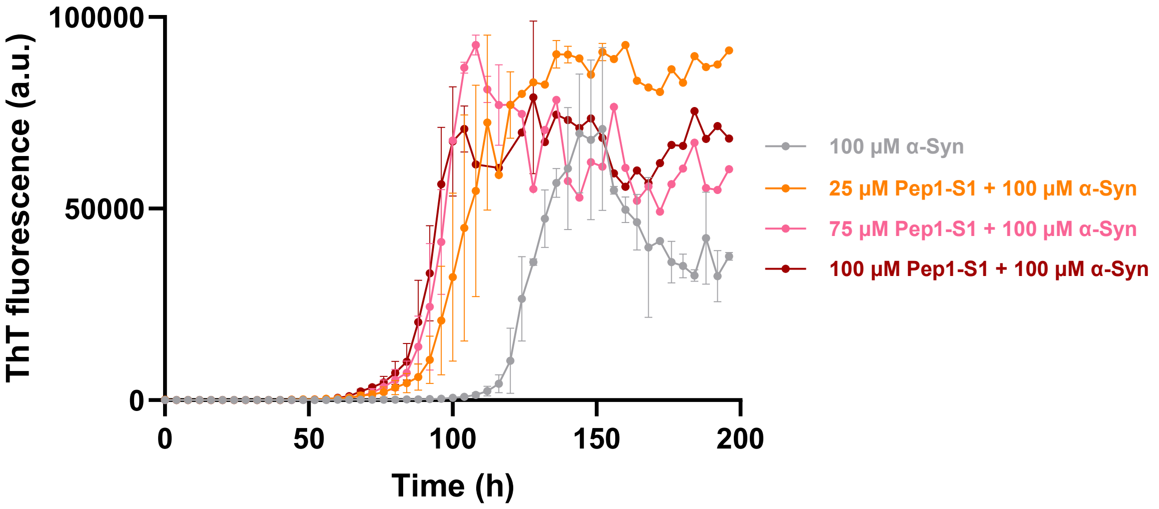


Figure S2. ThT fluorescence kinetics monitoring the aggregation of α-synuclein monomers (100 μM) in the absence or presence of Pep1-S1 peptide. Direct comparison shows the effects of 25, 75, and 100 μM Pep1-S1 on α-synuclein aggregation. Measurements were performed in triplicate (n = 3).

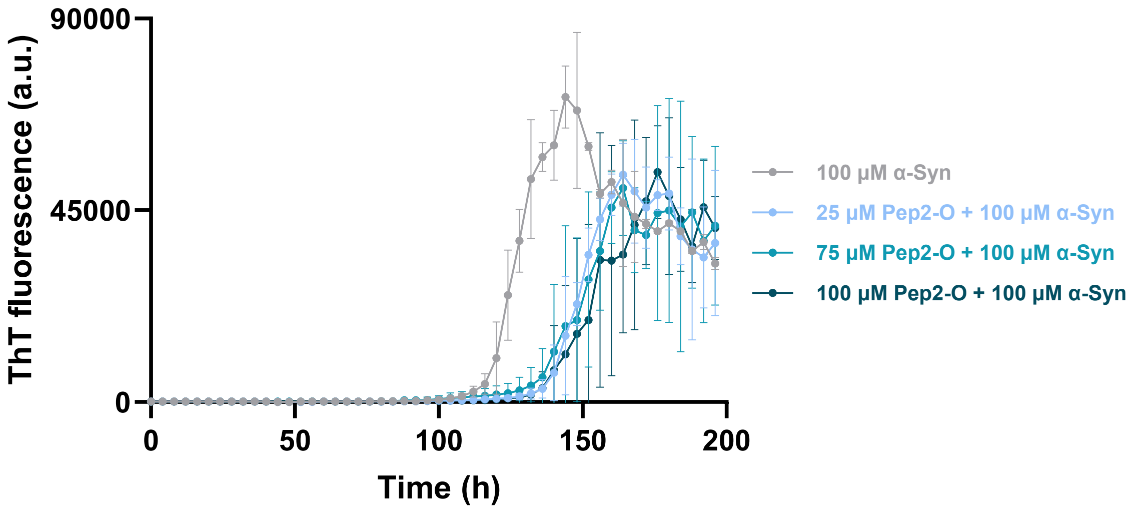


Figure S3. ThT fluorescence kinetics monitoring the aggregation of α-synuclein monomers (100 μM) in the absence or presence of Pep2-O peptide. Direct comparison shows the effects of 25, 75, and 100 μM Pep2-O on α-synuclein aggregation. Measurements were performed in triplicate (n = 3).

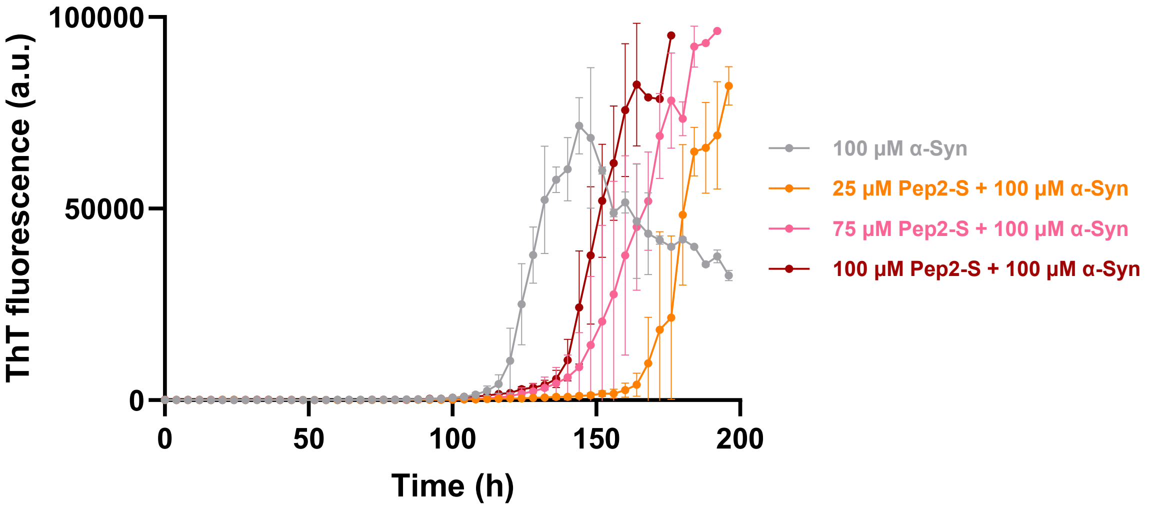


Figure S4. ThT fluorescence kinetics monitoring the aggregation of α-synuclein monomers (100 μM) in the absence or presence of Pep2-S peptide. Direct comparison shows the effects of 25, 75, and 100 μM Pep2-S on α-synuclein aggregation. Measurements were performed in triplicate (n = 3).

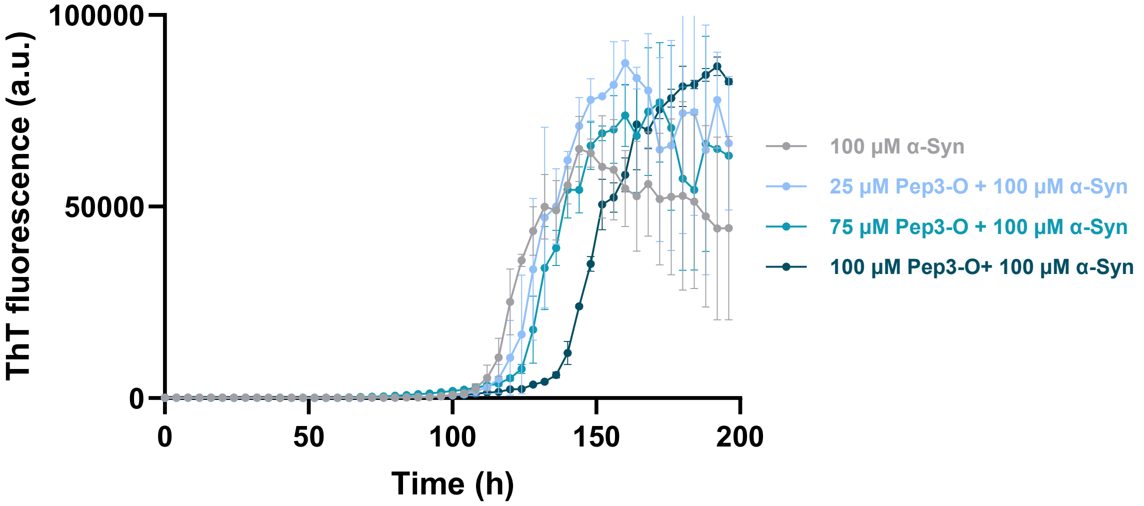


Figure S5. ThT fluorescence kinetics monitoring the aggregation of α-synuclein monomers (100 µM) in the absence or presence of Pep3-O peptide. Direct comparison shows the effects of 25, 75, and 100 µM Pep3-O on α-synuclein aggregation. Measurements were performed in triplicate (n = 3).

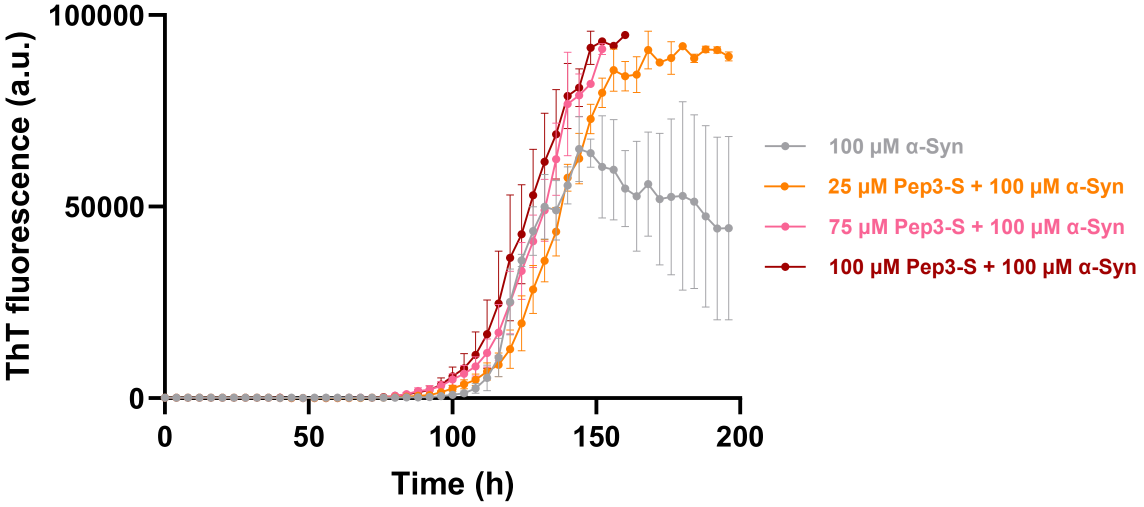


Figure S6. ThT fluorescence kinetics monitoring the aggregation of α-synuclein monomers (100 µM) in the absence or presence of Pep3-S peptide. Direct comparison shows the effects of 25, 75, and 100 µM Pep3-S on α-synuclein aggregation. Measurements were performed in triplicate (n = 3).

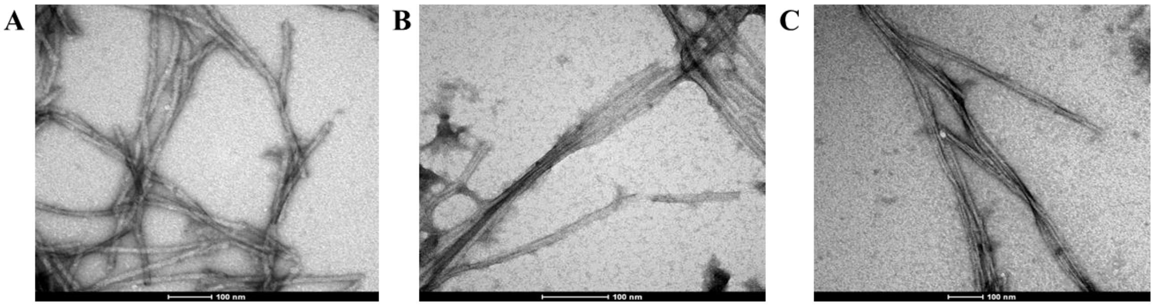


Figure S7. TEM of (A) 100 μM α-synuclein alone, 100 μM α-synuclein treated with (B) 25 μM Pep2-O or (C) 25 μM Pep2-S.

**General Synthesis.** Commercial chemicals were of reagent quality or better and were used without additional purification. Amino acids and resins were obtained from Ambeed, Inc (Arlington Heights, IL). All other reagents and solvents were obtained from Fisher Scientific and Sigma–Aldrich. Fmoc-Val-BTA and Fmoc-Gly-BTA were synthesized by the method of Mukherjee *et al.* (Mukherjee et al., 2015) Reactions were monitored by thin-layer chromatography with visualization by UV light or staining with $KMnO_4$. Chromatography was performed with columns of F60 Irregular Silica Gel, 40 - 63 µm (SiliCycle Inc, Québec City, Canada). Solvents and other volatile compounds are removed "under reduced pressure" using a rotating evaporator at water-aspirator pressure (<20 torr) and a 40°C water bath. Mass spectrometry was performed on either Agilent 6125 LC/MS or Agilent 6530 and 6546 Q-TOF.

**Peptide Synthesis.** Peptides were synthesized on a Liberty Blue 2.0 Microwave Peptide Synthesizer from CEM Corporation (Matthews, NC) using standard Fmoc-based methods starting from 2-Chlorotrityl chloride resin obtained from Chem-Impex International (Wood Dale, IL). Amide bond formation was achieved using 5 eq. of amino acid (0.2 M), 10 eq. of DIC (1 M), and 5 eq. of ethyl cyanohydroxyiminoacetate (1M) and heating to 60°C for 600 s. Fmoc deprotection was achieved with 20% *N*-methylpiperidine in DMF at a temperature of 75°C for 15 s, followed by heating to 90 °C for 50 s. For thioamide peptides, 1 equiv. of Fmoc-Val-BTA or Fmoc-Gly-BTA was manually coupled with 4 equiv. DIPEA for 1 h. Subsequent amide bond formation was achieved using 5 eq. of amino acid (0.2 M), 10 eq. of DIC (1 M), and 5 eq. of ethyl cyanohydroxyiminoacetate (1 M) and heating to 50°C for 600 s. Cleavage and global deprotection was achieved by a 3-hour treatment with a solution of 82.5% v/v TFA, 5% v/v phenol, 5% v/v water, 5% v/v thioanisole, and 2.5% v/v 1,2-ethanedithiol.

After evaporating the cleavage solution under a flow of nitrogen, the peptide was dissolved in 25% (v/v) MeCN in water. Peptides were purified by reversed-phase HPLC using an UltiMate™ 3000 Basic HPLC System from Thermo Fisher Scientific (Waltham, MA) on a Vydac 214TP 10µm C18 column from HiChrom (Wilmington, DE) using a linear gradient of 22–37% v/v B over 30 min (A: 0.1% v/v TFA in $H_2O$; B: 0.1% v/v TFA in MeCN) Subsequent lyophilization yielded white powders. Purified peptides were analyzed by LC/MS using Agilent ZORBAX Eclipse Plus C18 narrow bore column; 2.1 mm internal diameter; 50 mm length; 5-micron particle size; P.N. 959746-902 with a linear gradient of 5–95% v/v B over 15 min (A: 0.1% v/v formic acid in $H_2O$; B: 0.1% v/v formic acid in MeCN).

**Expression and purification of α-synuclein.** This method was performed as previously described for the expression and purification of α-synuclein. (Noh and Newberry, 2026)

**Turbidity assay.** The peptides were dissolved in 25 mM sodium phosphate buffer (pH 7.4), 0.1 M NaCl, and 1 mM LiOH. The corresponding concentrations were determined spectrophotometrically. (Anthis and Clore, 2013) Samples were prepared and loaded in triplicate into full area 96 well plates (100 µL per well),, then incubated at 37°C in a moist chamber under quiescent conditions with continuous shaking at 300 rpm. Data points were collected every 6 min. A wavelength of 600 nm was used to monitor amyloid fibril formation over time. The plates were read using a Synergy H1 microplate reader (BioTek, USA).

**Thioflavin T kinetic assay.** The corresponding concentrations of α-synuclein and short peptides were determined spectrophotometrically. (Anthis and Clore, 2013) 100 µM α-synuclein monomer alone or in the presence of short peptides was dissolved in 25 mM sodium phosphate buffer (pH 7.4), 0.1 M NaCl, and 25 µM ThT. Samples were prepared and loaded in triplicate into full area 96 well plates (100 µL per well), then incubated at 37°C in a moist chamber under quiescent conditions with continuous shaking at 300 rpm. Data points were collected every 5 min. The steady state ThT fluorescence was measured with excitation at 440 nm and emission at 485 nm. The plates were read using a Synergy H1 microplate reader (BioTek, USA).

**Transmission electron microscopy imaging.** Transmission electron microscopy (TEM) imaging was performed to visualize fibril morphology. Samples of α-synuclein, α-synuclein with 25 µM Pep2-O, and α-synuclein with 25 µM Pep2-S were collected from kinetic experiments and diluted 100-fold prior to imaging. A 5 µL aliquot of each sample was applied onto carbon-coated copper grids (200 mesh) and allowed to adsorb for 1–2 min. Excess liquid was blotted off with filter paper. Grids were optionally stained with 1% (w/v) uranyl acetate for 30–60 s for negative staining, followed by air drying. TEM images were acquired using a JEOL 1400 transmission electron microscope operating at 80–120 kV.

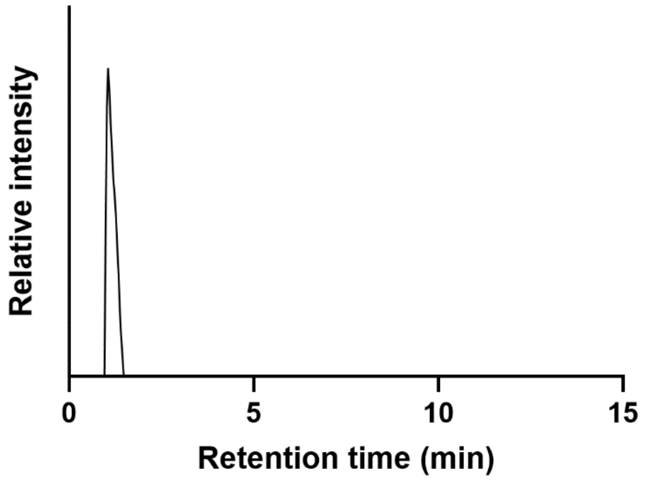


Analytical LC–MS trace of purified Pep1-O. Calculated monoisotopic exact mass for $[M+H]^+$ ($C_{26}H_{47}N_7O_9$): 601.7; found: 602.3.

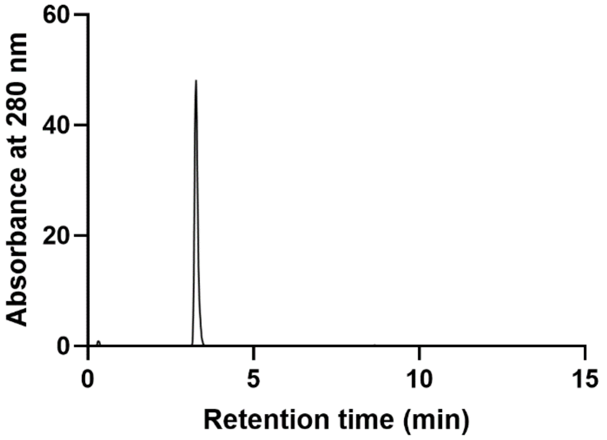


Analytical LC–MS trace of purified Pep1-S1. Calculated monoisotopic exact mass for $[M+H]^+$ ($C_{26}H_{47}N_7O_8S$): 617.8; found: 618.0.

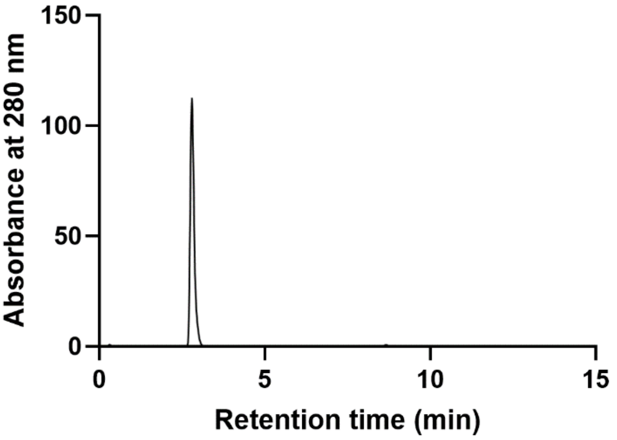


Analytical LC–MS trace of purified Pep1-S2. Calculated monoisotopic exact mass for $[M+H]^+$ ($C_{26}H_{47}N_7O_8S$): 617.8; found: 618.0.

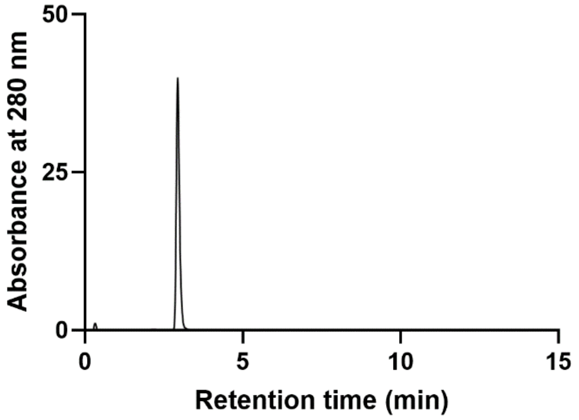


Analytical LC–MS trace of purified Pep1-S3. Calculated monoisotopic exact mass for $[M+H]^+$ ($C_{26}H_{47}N_7O_8S$): 617.8; found: 618.0.

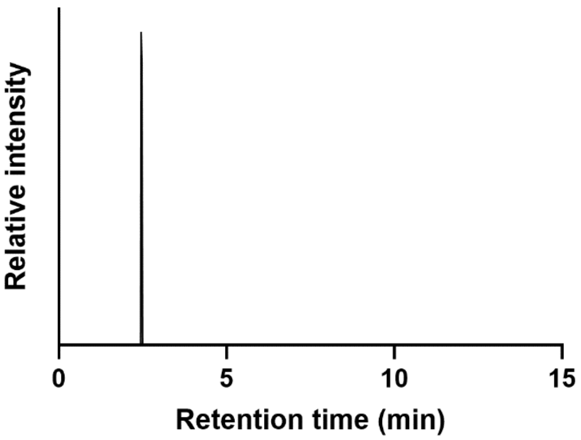


Analytical LC–MS trace of purified Pep2-O. Calculated monoisotopic exact mass for $[M+H]^+$ ($C_{27}H_{49}N_7O_9$): 615.7; found: 616.5.

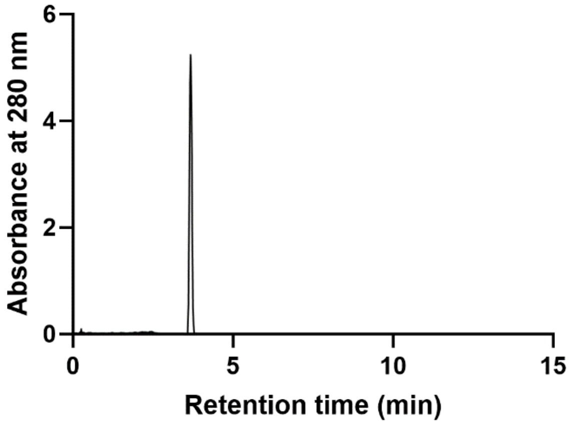


Analytical LC–MS trace of purified Pep2-S. Calculated monoisotopic exact mass for $[M+H]^+$ ($C_{27}H_{49}N_7O_8S$): 631.8; found: 632.5.

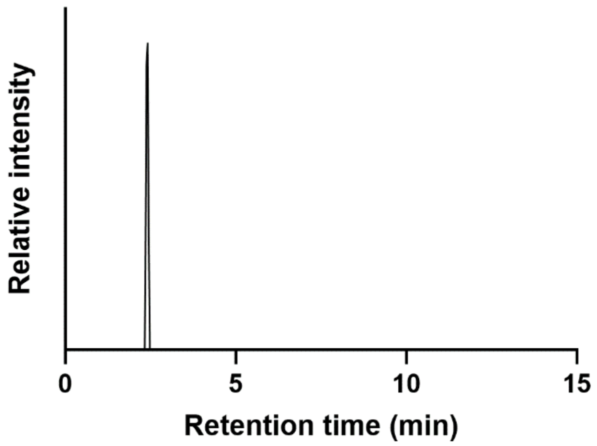


Analytical LC–MS trace of purified Pep3-O. Calculated monoisotopic exact mass for $[M+H]^+$ ($C_{27}H_{49}N_7O_9$): 615.7; found: 616.3.

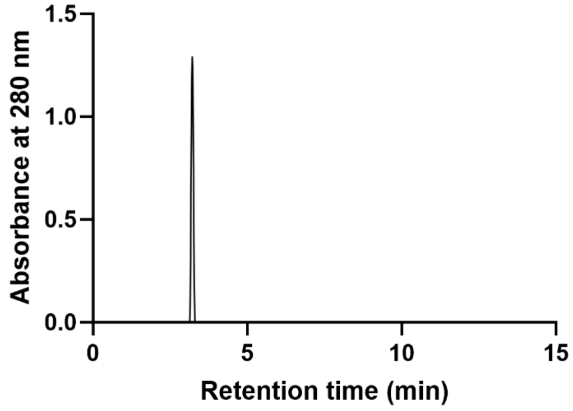


Analytical LC–MS trace of purified Pep3-S. Calculated monoisotopic exact mass for $[M+H]^+$ ($C_{27}H_{49}N_7O_8S$): 631.8; found: 632.3.

**References:**

Anthis NJ, Clore GM. Sequence-specific determination of protein and peptide concentrations by absorbance at 205 nm. *Protein Sci.* 2013; **22**(6): 851–858. https://doi.org/10.1002/pro.2253.

Mukherjee S, Verma H, Chatterjee J. Efficient Site-Specific Incorporation of Thioamides into Peptides on a Solid Support. *Org Lett.* 2015; **17**(12): 3150–3153. https://doi.org/10.1021/acs.orglett.5b01484.

Noh D, Newberry RW. Concentration-dependent mutational scanning probes the cellular folding landscape of α-synuclein in yeast. *Protein Sci.* 2026; **35**(2): e70456. https://doi.org/10.1002/pro.70456.
